# Transcriptomic and functional comparison of cells isolated from healthy and degenerated ovine intervertebral discs

**DOI:** 10.1101/2025.11.06.686904

**Authors:** Paul Humbert, Lucie Danet, Emmaëlle Carrot, Floriane Etienne, Boris Halgand, Frédéric Blanchard, Claire Vinatier, Jérôme Guicheux, Marion Fusellier, Catherine Le Visage, Romain Guiho

## Abstract

**Background:** Intervertebral disc degeneration (IVDD) is a leading cause of chronic low back pain and disability. Understanding the cellular and molecular mechanisms underlying disc degeneration is crucial for developing effective therapies. Sheep have emerged as a promising large-animal model for IVDD research due to their similarities with humans. They exhibit resembling spine anatomy and biomechanics, and they develop spontaneous age-associated degeneration of the disc. However, the specific cellular alterations occurring in annulus fibrosus (AF) and nucleus pulposus (NP) ovine cells during degeneration remain poorly characterized. In vitro, the benefits of using cells from aged sheep over young ones to mimic degenerative processes remain to be tested.

**Methods:** AF and NP cells from young and aged sheep were analysed using bulk RNA sequencing, with a focus on two hallmarks of IVDD: cellular senescence and metabolic alterations. Functional assays completed this focus by assessing cells response under basal conditions and after pro-degenerative stimuli (IL-1β, senescence induction). In addition, bulk transcriptomic data were deconvoluted using a reference single-cell RNA-seq dataset from healthy and degenerated human discs, and gene co-expression modules were compared across species.

**Results:** MRI and histological analyses revealed homogeneous mild degeneration across all lumbar discs in aged sheep, while lamb discs were uniformly healthy. Cells transcriptomic profiling identified robust age- and tissue-specific signatures, with aged NP and AF cells showing upregulation of inflammatory mediators, ECM-remodelling enzymes, and senescence-associated pathways. Cross-species analysis revealed shared transcriptional modules between aged sheep cells and human degenerated disc cells, supporting the translational relevance of the ovine model. Remarkably, young and aged cells shared a similar functional behaviour when exposed to stress-related stimuli.

**Conclusions:** This work confirms the compatibility of sheep cells with in vitro testing and their relevance to model human IVDD. Cross-validation with human single-cell data further highlights common pathogenic pathways, reinforcing the translational potential of the model. However, no added benefits were found in using older animals compared to younger ones as cell sources in functional assays.

**Highlights:**

- Transcriptomic profiling of AF and NP cells from young and aged sheep
- Aged cells show inflammatory, ECM-remodelling and senescence signatures
- Deconvolution with human scRNA-seq links aged ovine and degenerated discs
- Sheep cells retain in vitro responsiveness to pro-degenerative stimuli
- Supports the ovine model as a translational tool for IVDD research

## 1. Introduction

Chronic low back pain (LBP) is the leading cause of years lived with disability and affects over 600 million patients worldwide [1]. This debilitating condition is particularly prevalent in the aging population and among individuals with decreased levels of physical activity. Therefore, it is expected to affect an increasing number of people in the near future, with a considerable burden, both from a societal and individual patient perspective [2]. The urgent need for better management of LBP is one of the World Health Organization (WHO) priorities, which is reflected by the implementation of the Decade of Healthy Aging for 2021-2030. Chronic LBP has multiples causes, but among them, intervertebral disc degeneration (IVDD) is known to be the major contributor [3].

The intervertebral disc (IVD) is composed of two main regions - the annulus fibrosus (AF), a fibrous outer ring, and the nucleus pulposus (NP), a gel-like core – connected to adjacent vertebrae by cartilaginous endplates (CEP). Each of those areas contains specialized cells responsible for matrix secretion but the NP also has a contingent of cells from notochordal origin. IVDD is characterized by a global loss of cellularity, cellular senescence, extracellular matrix (ECM) alterations and dehydration, increased expression of MMPs and inflammatory factors, leading to a loss of IVD biomechanical properties, ectopic ingrowth of sensory nerves and blood vessels, pain and disability. Clinical management, including anti-inflammatory and analgesic drugs for early degenerative stages, and surgery for advanced ones, only aims at controlling pain but does not address the underlying pathological processes [4]. Given the lack of curative options and the urgent need to identify and test new therapeutic opportunities, the development of relevant preclinical models that faithfully recapitulate human IVDD is essential.

Among laboratory animals for IVDD research, the sheep has emerged as a promising model organism [5–7] due to anatomical similarities with the human spine [8,9] (e.g., comparable disc size and biomechanical constraints) and spontaneous radiological changes observed in aging animals resembling those found in human patients [10]. This natural degeneration of sheep IVD may be caused by a decline in the notochordal cell population in adult animals [11], as reported in human physiology [12]. The ovine model of IVDD has the benefits of allowing clinically relevant surgical procedures [13], the generation of whole disc ex vivo explants [14] and primary cultures of isolated cells [15]. This wide range of applications is particularly interesting for translational research. However, despite numerous preclinical and in vitro studies using this animal model, it should be stressed that ovine disc cells and their behaviour in culture remain poorly characterized.

This study explores the relevance of cellular models derived from AF and NP tissues of both lambs and aged sheep, aiming to characterize their behaviour in vitro and gain insights into the age-related disc degeneration processes. To this end, we performed an in-depth transcriptomic analysis of expanded cells isolated from AF and NP in lambs and sheep to identify key differences related to the age of the donor animal or the tissue of origin, with a focus on cellular senescence and energy metabolism. First, cellular senescence has emerged as a hallmark of IVDD, contributing to matrix breakdown and the secretion of pro-inflammatory mediators [16]. Second, an imbalance in energy metabolism between oxidative phosphorylation (OXPHOS) and glycolysis has also been recently identified as a potential driver of disc cell dysfunction, and its alteration could be critical to understand the pathophysiology of IVDD in the years to come [17]. In recent years, the advent of transcriptional studies on single cells has enabled the scientific community to compile important datasets to evaluate the heterogeneity of cellular states in healthy and diseased human discs. To compare the transcriptomes of our ovine cells with those of human discs, we cross-examined the dataset generated by Cherif et al. [18], a single-cell RNAseq study of NP and AF from a healthy and a diseased human disc. Ultimately, using relevant functional assays, we compared NP cells derived from aged sheep with those from young lambs, under basal conditions and after pro-degenerative challenges such as IL-1β exposure or senescence induction.

## 2. Methods

### 2.1. Sample Collection

Lumbar IVDs were collected from four 6-month-old female lambs and four female sheep between 7-8 years-old (Vendée breed, GAEC HEAS, Les Rabelais, Ligné, France) in the accredited Centre of Research and Pre-Clinical Investigations (ONIRIS, National Veterinary School of Nantes). The sheep received a bolus hypocoagulant dose of heparin (50 IU/kg) and were euthanized with an overdose of barbiturates (60 mg/kg) following the best practices and legislation (European Directive 2010/63/EU). Post-euthanasia, MRI of the lumbar spine was performed to obtain sagittal T2-weighted images (TE: 86 ms, TR: 3,000 ms; slice thickness: 3 mm), using a 1.5 T MRI scanner (Magnetom Essenza, Siemens Medical Solutions). OsiriX 9 software (Osirix Foundation) was used to analyse MRI images, and each lumbar IVD was scored using the Pfirrmann grading system [19]. Under aseptic conditions, the lumbar spines were extracted via a posterior approach, and the IVDs were individualized with part of the vertebral bodies using an oscillating saw. The sections were then transported to the cell culture room in sterile media [14].

### 2.2. Histological analysis

One representative lumbar IVD per animal was processed for histological analysis, after fixation with Paraformaldehyde (PFA, Sigma-Aldrich) 4% for 5 days, followed by decalcification (Shandon TBD-2 Decalcifier, Thermo Fisher Scientific) for 3 weeks and a post-fixation step, after abundant rinsing, in 4% PFA for 24 hours. IVDs were then frozen in isopentane cooled with dry ice and embedded in SCEM medium (Section Lab). Coronal cryosections of 10 µm thickness were made with a cryostat (CryoStar NX70, Thermo Fischer Scientific) and stained with either hematoxylin-eosin-saffron (HES) or Alcian blue, according to standard procedures. Images of the stained sections were acquired with a slide scanner (NanoZoomer, Hamamatsu Photonics) and visualized with NDP.view2 software (Hamamatsu Photonics).

### 2.3. Cell isolation and culture

AF and NP cells were isolated as previously described [20]. Briefly, the 5-remaining lumbar IVDs were dissected from the vertebral bodies, AF and NP were separated and diced into pieces. Tissue fragments were rinsed three times in HBSS (L0606, Biowest) with 2% penicillin/streptomycin (P/S; 15070-063, Gibco) for 2 minutes. The extracellular matrix was then digested with 0.05% hyaluronidase (H4272, Sigma-Aldrich) in HBSS at 37 °C for 15 minutes. After two rinses with HBSS, 15–20 mL of 0.2% trypsin (T9935, Sigma-Aldrich) in HBSS were added to each sample and incubated at 37 °C for 30 minutes. After two additional rinses with HBSS, 15–20 mL of 0.25% collagenase (C5138, Sigma Aldrich) were added in complete culture medium, i.e. DMEM high glucose and pyruvate (31966-021, Gibco) supplemented with 10% foetal bovine serum (FBS eurobio scientific CVFSCF00-01, Lot n° S76214), 1% P/S, and 0.1% Amphotericin B (15290, Gibco), and the mixtures were incubated at 37 °C overnight. The recovered suspension was filtered through a 70-μm-pore filter and centrifuged for 5 minutes at 300 g. The cells were counted and seeded at 10 000 cells /cm² in T75 flasks (Sarstedt). AF and NP cells were grown at 37 °C and 5% CO_2_.

### 2.4. RNA sequencing and transcriptomic analysis

Total RNA was extracted from lamb and sheep AF and NP cells at passage 1 (P1) using NucleoSpin RNA XS kit (Macherey-Nagel, 740902). Library preparation was performed with Illumina® Stranded mRNA Prep kit. Sequencing was then performed on an Illumina HiSeq with NovaSeq 6000 SP Reagent Kit (200 cycles) v1.5. Sequencing data was then aligned with STAR using Ovis_aries_rambouillet.ARS-UI_Ramb_v2.0.111 genome. Secondary analysis was performed using {R 4.3.3} on RStudio (2023.06.0+421). Differential expression was performed using {DESeq2 1.38.1}. Geneset Enrichment Analysis (GSEA) was performed using the GSEA software (v4.3.2 - UC San Diego and Broad Institute). Gene sets used in this study were either obtained from public databases (gene ontology (GO), KEGG, REACTOME, molecular signatures database - Broad Institute) or published literature: senescence profiling [21] and Senescence-Associated Secretory Phenotype (SASP) [22]. Weighted Gene Co-Expression Network Analysis (WGCNA) was performed using {WGCNA 1.72} R package to identify pathways and networks associated with aging. A public dataset of single-cell RNAseq on human disc cells (GSE199866) [18] has been cross-analysed using a {Seurat 5.3.0} R pipeline. Briefly, after quality control (filtering on high mitochondrial genes percentage and low number of features, doublet removing with {DoubletFinder 2.0.4}), integration was performed with {rliger 2.2.0} using k = 20 and lambda = 40 settings. Clusters were annotated based on the literature consensus [23–25]. Bulk deconvolution based on scRNA-seq was performed with {MuSiC 1.0.0} [26]. scRNA-seq single sample GSEA (ssGSEA) based on WGCNA module score was performed using {escape 2.4.0}.

### 2.5. Cellular senescence characterisation

NP cells at P3 were seeded in 96-well plate (Sarstedt) at 5,000 cells per well and either: treated with recombinant ovine IL-1β (5 ng/mL; QP6215-YE, enQuireBio) for one week with treatment renewal every 3 days; treated with etoposide (10 μM; E1383, Sigma-Aldrich) for 24 hours then maintained in culture for one week; or serum-starved for one week, as previously described [27]. Senescence-associated (SA)-β-galactosidase and 5-ethynyl-2′-deoxyuridine (EdU) incorporation staining were performed on the same cells. Briefly, at day 7, EdU (10 µM) was added to the culture medium overnight. SA-β-galactosidase staining was performed at day 8 using the Senescence Cells Histochemical Staining Kit (CS0030, Sigma-Aldrich) following the manufacturer’s instructions. EdU incorporation staining was performed afterward using Click-iT EdU Cell Proliferation Kit (Invitrogen) following the manufacturer’s instructions. DAPI was used as a nuclear counterstain. Imaging was performed on a Zeiss Macroscope colour fluorescence, using Zen Blue software v3.7. Analysis was performed with CellProfiler software v4.2.5.

### 2.6. Measurement of mitochondrial respiration

NP cells at P3 were seeded in duplicate in XF96 plates at 60,000 cells per well and treated with recombinant ovine IL-1β (1 ng/mL) or serum-starved. After 24 hours, the culture medium was replaced with DMEM phenol red-free medium pH 7.4 (5030, Sigma-Aldrich), supplemented with glutamine (4 mM), glucose (25 mM), and pyruvate (1 mM). The plate was incubated for 45 minutes at 37°C without CO2 to degas and allow accurate measurements. For the mitostress assay, oxygen consumption rate (OCR) measurements were performed every 5 minutes before and after the successive injections of oligomycin (2 µM), carbonylcyanide-3-chlorophenylhydrazon (CCCP, 4 µM) and a 1:1 mixture of rotenone/antimycine A (1 µM each) using the Seahorse XF Pro analyser (Agilent). Data analysis was performed using Wave software (Agilent) as previously described [28].

### 2.7. Statistical analysis

All the results are presented as means ± SEM. When applicable, points on graphs represent the values for the different biological replicates. The statistical analyses were performed using GraphPad Prism software v10.0. Statistical significance was determined using the appropriate test, detailed in each figure legend.

## 3. Results

### 3.1. Sheep develop spontaneous mild intervertebral disc degeneration with age

As the first step of our experimental process (**Figure 1A**), eight animals were obtained from the same breeder: four female sheep between 7-8 years old and 4 sex-matched female lambs. Six-month-old lambs exhibited healthy IVDs based on MRI scoring, with a homogeneous Pfirrmann score of 1 for all discs scored. In contrast, aged sheep showed an increased score around 2, suggesting a homogeneous degenerative profile at the lumbar level (**Figure 1B-C**). All animals had six lumbar IVDs. Of note, one lamb presented an L5 vertebral deformity, which did not affect the Pfirrmann score or tissue processing. MRI-based observations were confirmed with histological sections and staining for 1 disc per animal, selected as representative of the mean degeneration score for the animal (**Figure 1D**). Consistent with our previous report [10], healthy discs showed higher cellularity, while "ghost cells” (i.e. residual space in the matrix after putative cell death) were seen mostly in mildly degenerated IVDs (**Figure 1E**). The presence of vertebral growth plates and the overall larger size of young discs illustrate that the lambs were not skeletally mature (**Supplementary Figure S1A**).

**Figure 1.**
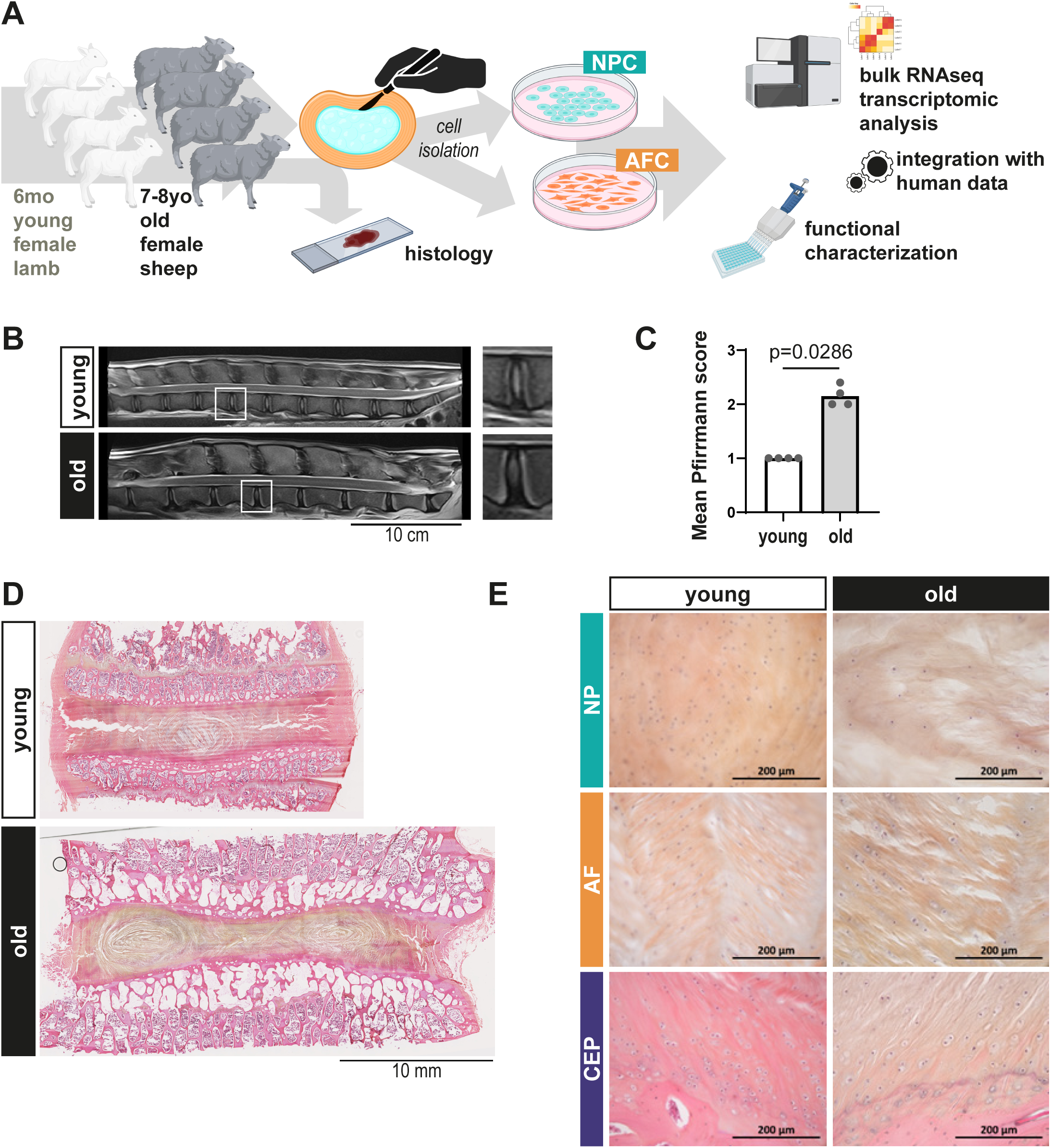
Sheep develop spontaneous mild intervertebral disc degeneration during aging. **A.** Study design. Lumbar IVDs were collected from 4 female lambs (6-month-old) and 4 female sheep (7-8 years old). Medullar MRI of the thoracolumbar sheep spines were graded using the Pfirrmann system. One representative IVD per animal was processed for histology, the 5 others were dissected for AF and NP cells, separate isolation. Bulk RNA sequencing was performed for transcriptomic analysis. Functional characterization was then carried out. **B.** Representative MRI images and **C.** mean lumbar IVDs Pfirrmann score per animal. Scale bar, 10 cm. p-values were calculated using the Mann-Whitney test (N=4 animals per group, n=6 IVDs scored per animal). **D.** Representative hematoxylin-eosin-saffron (HES) staining images of all discs and **E.** magnifications of areas of interest. Scale bars, D: 10 mm, E: 200 μm*. AF: Annulus Fibrosus, NP: Nucleus Pulposus, CEP: Cartilaginous Endplate*.

Cells could be efficiently isolated from both AF and NP tissues for all animals (**Supplementary Figure S1B-D**). More NP cells (NPC) tended to be retrieved (**Supplementary Figure S1C**), NPC from old animals exhibited a slightly longer doubling time compared to age-matched AF cells (AFC) (**Supplementary Figure S1D**). Overall, primary cultures from both tissues and age groups exhibited comparable growth behaviour under standard culture conditions and a similar morphology (**Supplementary Figure S1B**).

### 3.2. Disc cells from animals of different ages and from different tissues maintain a specific transcriptomic profile in culture

We next performed a bulk RNA sequencing on 16 cell samples at P1 (8 biological replicates isolated from either NP or AF tissues, including 4 young and 4 old animals). As the breed of sheep used in this study was the *Vendée* breed, the sequencing results were aligned with the genome of a neighbouring breed available on Ensembl: the Rambouillet sheep (*Ovis aries rambouillet*). 30,722 automatically identified genes and pseudo-genes were used for alignment, 12,159 were kept in this study after low count genes pre-filtering. 91.2 % were annotated using NCBI Wikigene. Forty-eight (0.4%) were manually curated using human orthologous sequences (**Supplementary Figure S2**). The first observation following this whole transcriptome analysis was the significant heterogeneity of our different biological samples, a heterogeneity that was particularly pronounced in older animals and in AF tissues, as shown by the dispersion and the absence of clear clustering on the PCA plot (**Figure 2A**). Although highly heterogeneous, disc cells from animals of different ages nevertheless retained an age-specific profile in culture (Figure 2B). Differential Gene Expression between cells isolated from young and old animals identified 248 differentially expressed genes (DEGs) for NPC (**Supplementary Table 1**) and 455 DEGs for AFC (**Supplementary Table 2**). Some of this DEGs are commonly upregulated in AFC and NPC isolated from old animals, including *GREB1*, *ESR1* linked to estrogen-response or *EEPD1*, *TDP2*, and *AKR7A2* potentially involved in DNA damage-response and stress-response. On the other side of the volcano plots, developmental factors *SOX6* and *NCAM1* are overexpressed in both NPC and AFC isolated from young lambs, alongside *BASP1*, a known marker of juvenile human discs [29]. The whole transcriptome analysis also provides an overall view of the genes encoding collagens, revealing that their transcriptomic signature can distinguish not only cells isolated from AF or NP, but also AFC originating from young or old animals (**Figure 2C**). Notably, for NPC, the age-separation based solely on collagens was not as clear-cut. Interestingly, the genes encoding matrix proteins, previously highlighted in a proteomic study of aging human discs [30], helped distinguish NPC from AFC at any age, although with significant inter-sample heterogeneity (**Supplementary Figure S3A**).

**Figure 2.**
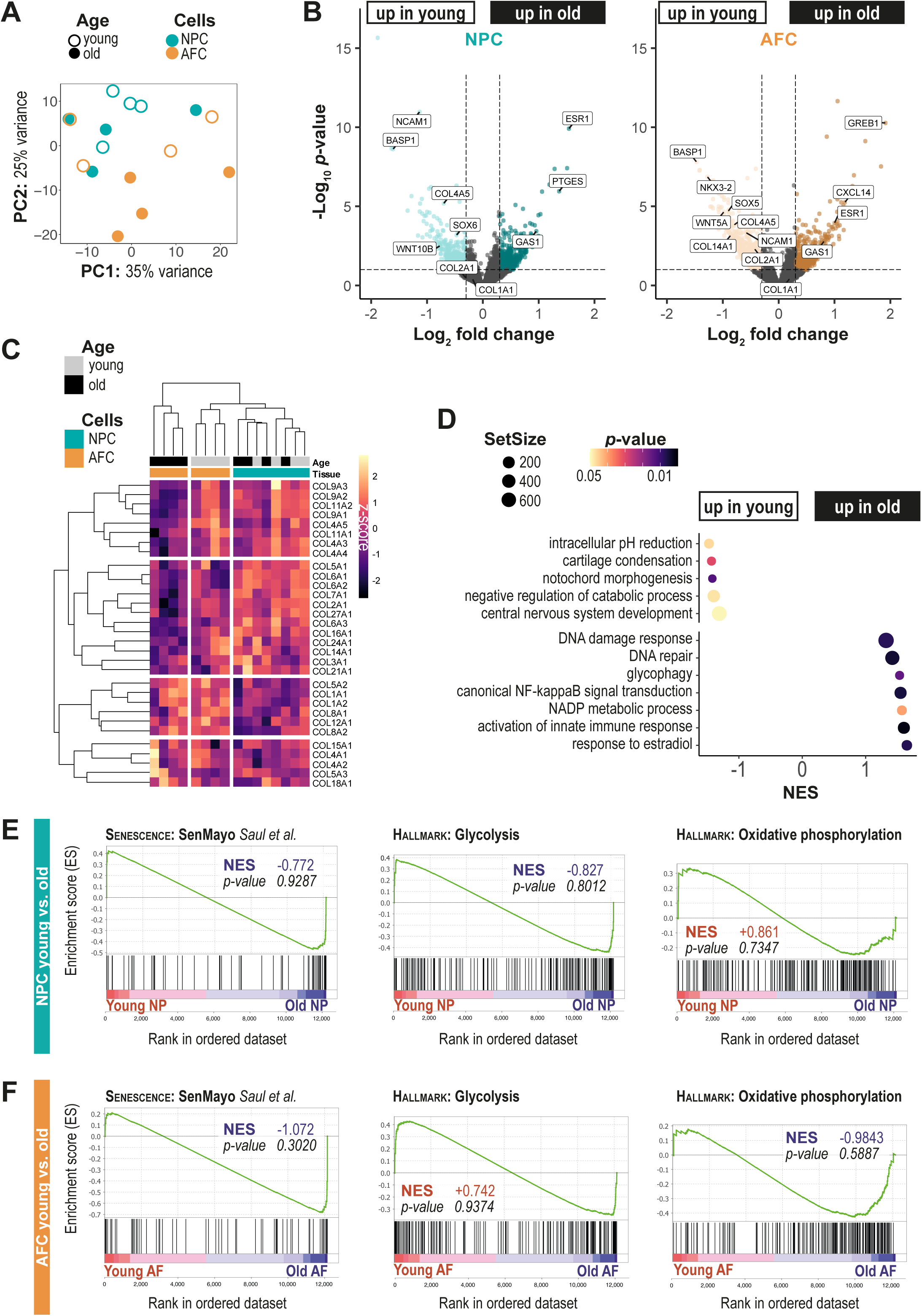
Transcriptomic Profiling and Differential Gene Expression in Ovine Disc Cells from Young and Old Animals. **A.** Principal component analysis (PCA) of cells transcriptomes obtained from old (empty dot) and young (plain dot) animals, and from AFC (orange) and NPC (turquoise). **B.** Volcano plot of differential gene expression in NPC (left) or AFC (right) from young versus old animals using the Wald test. **C.** Clustering on collagens transcripts expression **D.** Geneset Enrichment Analysis (GSEA) young versus old samples using the permutation test. **E.** GSEA enrichment plots for 3 genesets relative to cellular senescence (*SenMayo*) and energy metabolism (glycolysis and OXPHOS) for young NPC versus old NPC, and **F.** for young AFC versus old AFC. *NES: normalized enrichment score*.

To complete the unbiased transcriptomic analysis of our cultured ovine disc cells, we performed a pathway enrichment analysis between cells isolated from young animals vs. old animals (**Figure 2D**). Notable differences were observed in pathways related to development (e.g., cartilage development, notochord and central nervous system development), which were significantly enriched in young samples. In contrast, cellular stress and inflammation pathways (e.g., DNA repair, NF-κB activation, innate immune response activation) were enriched in old samples. Moreover, a switch in metabolism-related processes (e.g., glycophagy and NADP metabolic process in old samples, regulation of intracellular pH in young samples) was observed (**Figure 2D**).

In order to verify whether predicted cellular senescence and energy metabolism imbalance described in IVDD could be confirmed in cells from aged animals compared to young ones, GSEA was performed separately on NPC and AFC using 3 relevant geneset: the *SenMayo* geneset [21] for senescence and both *hallmark*:glycolysis and *hallmark*:OXPHOS for energy metabolism imbalance. If Normalized Enrichment Scores (NES) seemed to indicate indeed an enrichment for cellular senescence processes in NPC and AFC from old animals (negative scores in favour of the “old” condition, respectively NES=-0.772 and NES=-1.072 - **Figure 2E,F**) and a metabolic switch towards glycolysis at the expense of OXPHOS in NPC from old animals (negative glycolysis NES and positive OXPHOS NES - **Figure 2E**), p-values did not reach significance.

On the tissue of origin aspect, disc cells from animals of both ages retained also a tissue-specific core profile (**Supplementary Figure S3B, C**). 253 DEGs have been identified between AFC and NPC from young animals (**Supplementary Table 3**). Among these DEGs, the genes encoding for the transcription factors *FOXP2* was overexpressed in AFC (Foldchange > 2) and *RUNX2* was overexpressed in NPC (Foldchange > 2) (**Supplementary Figure S3B**). Surprisingly, the genes recognised to be markers differentiating NP and AF tissues, namely *ACAN*, *COL2A1*, and *PAX1* expressed by NPC, and *COL1A1* and *HTRA1* expressed by AFC, failed to reach the significance threshold. *CA12*, a proposed marker of NPC was a notable exception with a significant over expression in NPC from young animals. This contrasts with samples from older animals, where these transcriptomic markers are clearly discriminating AFC from NPC (Foldchange > 1.3, 1.5, 1.4, 1.6 for *ACAN*, *COL2A1*, *PAX1*, *CA12* in NPC and >1.6, 1.4 for *COL1A1*, *HTRA1* in AFC) (**Supplementary Figure S3C**). Overall, transcriptomic differences were more importantly marked in cell populations derived from aged animals, with 1117 DEGs between AFC and NPC (**Supplementary Table 4**). We then performed pathway enrichment analysis between ovine NPC and AFC (**Supplementary Figure S3D**), which highlighted differences in the cellular processes involved (e.g., glucose response and acyl-CoA biosynthesis for AFC; glycosylation and tryptophan metabolism for NPC). These differences may reflect NPC and AFC different adaptation strategies to the in vitro environment.

To further explain the relatively modest number of DEGs between NPC and AFC transcriptomes, particularly in young animals, we took advantage of the human scRNA-seq dataset GSE199866 [18]. In line with our observations, the UMAP representation revealed a strong similarity between the transcriptomes of human cells freshly isolated from AF and NP. These cell types clustered together according to their cellular states rather than being separated by tissue of origin (**Supplementary Figure S3E**).

Altogether, our data highlighted that ovine cells isolated and cultured from animals of different ages and from distinct tissues maintained an age- and tissue-specific transcriptomic signature.

### 3.3. Cells derived from aged ovine discs exhibit human disc degeneration signatures

To further dissect the intra-samples heterogeneity observed in our transcriptomic dataset, we combined two complementary approaches: bulk RNA-seq deconvolution using a single-cell reference, and co-expression network analysis via Weighted Gene Co-Expression Network Analysis (WGCNA).

In order to identify the sub-populations of NPC and AFC present in our disc ovine culture, we first use bulk deconvolution technique based on known reference cell annotation (**Figure 3A**). For this purpose, the scRNA-seq GSE199866 dataset was used as previously. Cell clusters were annotated (**Figure 3B, Supplementary Figure S4A**) and their transcriptomic profile used as a reference to infer cellular states present in each ovine sample (**Figure 3C**). Interestingly, we highlighted that in vitro culture of disc cells led to a marked enrichment in fibrotic (65–75% in AF, ∼90% in NP) and progenitor-like cell states, regardless of donor age (**Figure 3C**). Notably, these two cell states represented minor populations in the native human disc (respectively 2% and 5% of the disc cells) and were not typically associated with IVDD (**Supplementary Figure S4B**).

**Figure 3.**
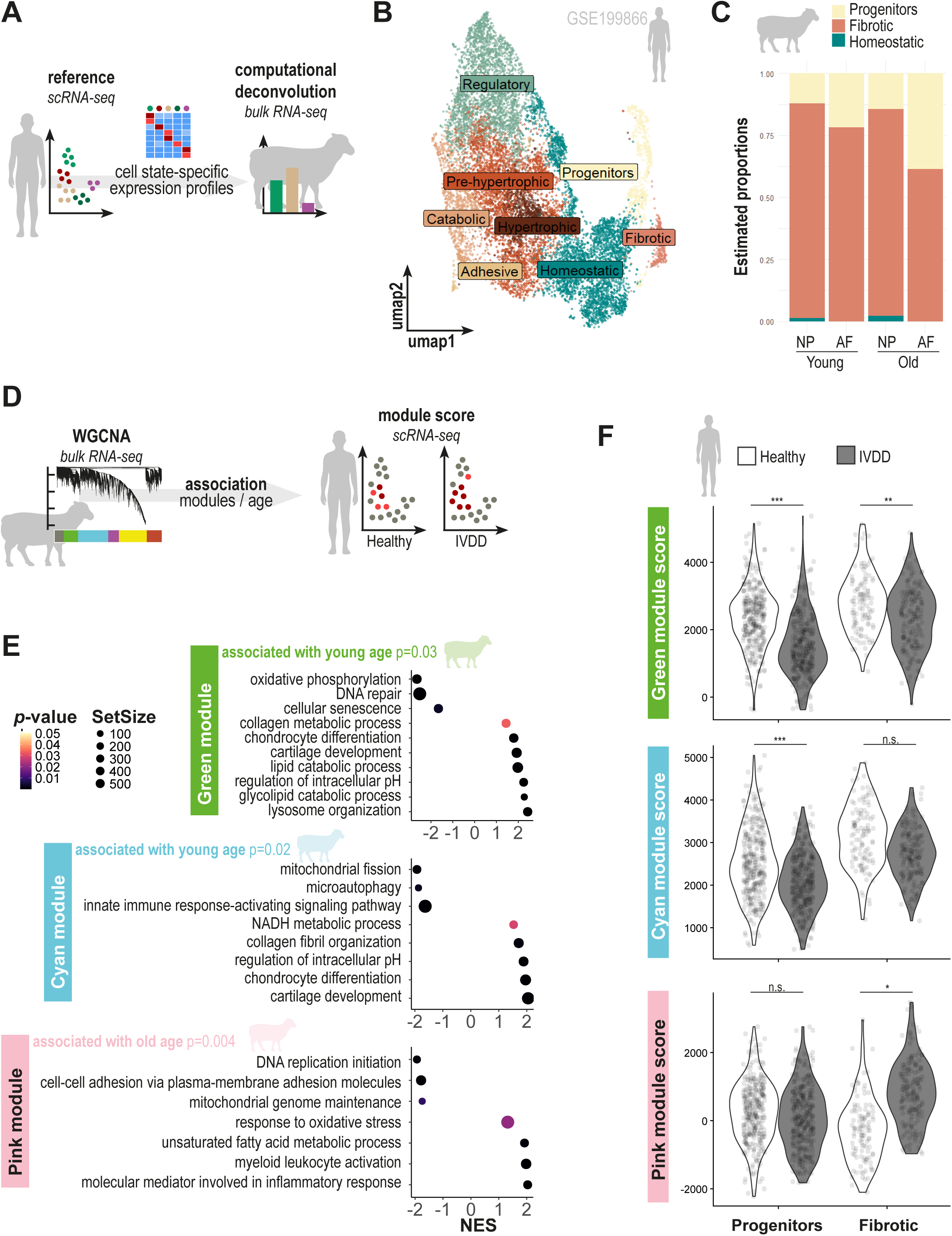
Cells from ovine aged disc express human disc degeneration signatures. **A.** Ovine bulk RNA-seq deconvolution approach based on a human scRNA-seq reference. The expression profiles of annotated cellular states served as a basis for computational deconvolution. **B.** UMAP of scRNA-seq human dataset (GSE199866) combining AFC and NPC transcriptome from one healthy and one IVDD intervertebral disc. Cellular states were annotated using literature consensus. **C.** Estimated proportion of cellular states in ovine samples by MuSiC deconvolution strategy. **D.** Bioinformatic approach to the weighted correlation network analysis (WGCNA) enabling the identification of gene modules associated with the age of ovine IVD. The scores associated with these modules were calculated for healthy and degenerated human IVD cells. **E.** Ranked GSEA of the genes associated with *Cyan* and *Green* (up in young animals) and *Pink* (up in old animals) modules. **F.** Top-gene modules used as ssGSEA score on human progenitor and fibrotic cells from healthy (clear) and IVDD (grey) disc. p-values were calculated with Wilcoxon signed-rank test, ns: not significant, *:p<0.05; **:p<0.01; ***:p<0.001.

To explore underlying gene expression programs, we applied WGCNA across all 16 bulk RNA-seq samples (**Figure 3D**) and identified six co-expression modules (**Supplementary Figure S4C**). Besides gene modules associated with AF tissue (*Yellow* and *Magenta* modules) or NP tissue (*Tan* and *Brown* modules), WGCNA identified 3 modules associated with young (*Cyan* and *Green* modules) or old age (*Pink* module) (**Supplementary Figure S4D**). For subsequent analysis, we focused on these 3 modules of interest. Functional annotation of these modules highlighted key biological processes differentially active with aging (**Figure 3E**). The *Green* module was positively enriched for pathways related to cartilage development and negatively enriched for DNA repair and cellular senescence. The *Cyan* module showed enrichment in chondrocyte differentiation and NADH metabolic processes, while being negatively associated with microautophagy, mitochondrial fission, and innate immune signalling. In contrast, the *Pink* module, upregulated in aged samples, was enriched for myeloid cell activation, oxidative stress response, and unsaturated fatty acid metabolism, while being negatively associated with DNA replication initiation and mitochondrial genome maintenance. These distinct co-expression signatures suggest that aging is accompanied by a coordinated shift away from anabolic and developmental programs toward inflammatory, metabolic, and stress-related responses, consistent with hallmarks of disc degeneration.

To assess the relevance of these ovine transcriptional programs to human IVDD, we projected the top 30 hub genes from each module onto the human scRNA-seq dataset (**Figure 3F, Supplementary Table 5**). Using single-sample GSEA (ssGSEA), we computed enrichment scores for each module across individual fibrotic and progenitor cells, two states commonly observed in vitro according to the bulk deconvolution. This analysis revealed a significantly higher proportion of cells with high *Green* and *Cyan* scores in healthy human discs compared to IVDD samples. Conversely, cells with high *Pink* module scores were significantly more frequent in fibrotic cells from degenerated discs (**Figure 3F**). These results indicate that the transcriptional modules enriched in young ovine cells reflect a healthy disc state, while those upregulated with age align with molecular signatures of human disc degeneration.

### 3.4. NPC from young and aged ovine disc share similar senescence and energy metabolic responses to stress-related stimuli

Based on the observed discrepancy in enriched pathways related to inflammation, senescence and energy metabolism, we sought to better characterize cellular functions associated with these processes. Given their central position in the pathophysiology of IVDD, the role of NPC in disc degeneration has been the subject of numerous studies in the literature. In this section, we chose to functionally explore the stress-response of these cells, at baseline and after stimuli. Cell proliferation, SA-β-Gal expression and mitochondrial respiratory profiles of NPC from young and old animals in response to various stimuli were evaluated (**Figure 4**, **Supplementary Figure S5**). IL-1β treatment mimicked inflammatory processes, etoposide treatment served as an efficient senescence inducer, and serum starvation was used as a basic culture condition known for influencing cell proliferation and metabolism. To validate the responsiveness of NPC to IL-1β, quantification of *MMP3* and *IL6* transcript expression was performed, showing significant overexpression of these markers in cells from both young and aged animals (**Supplementary Figure S5A**).

**Figure 4.**
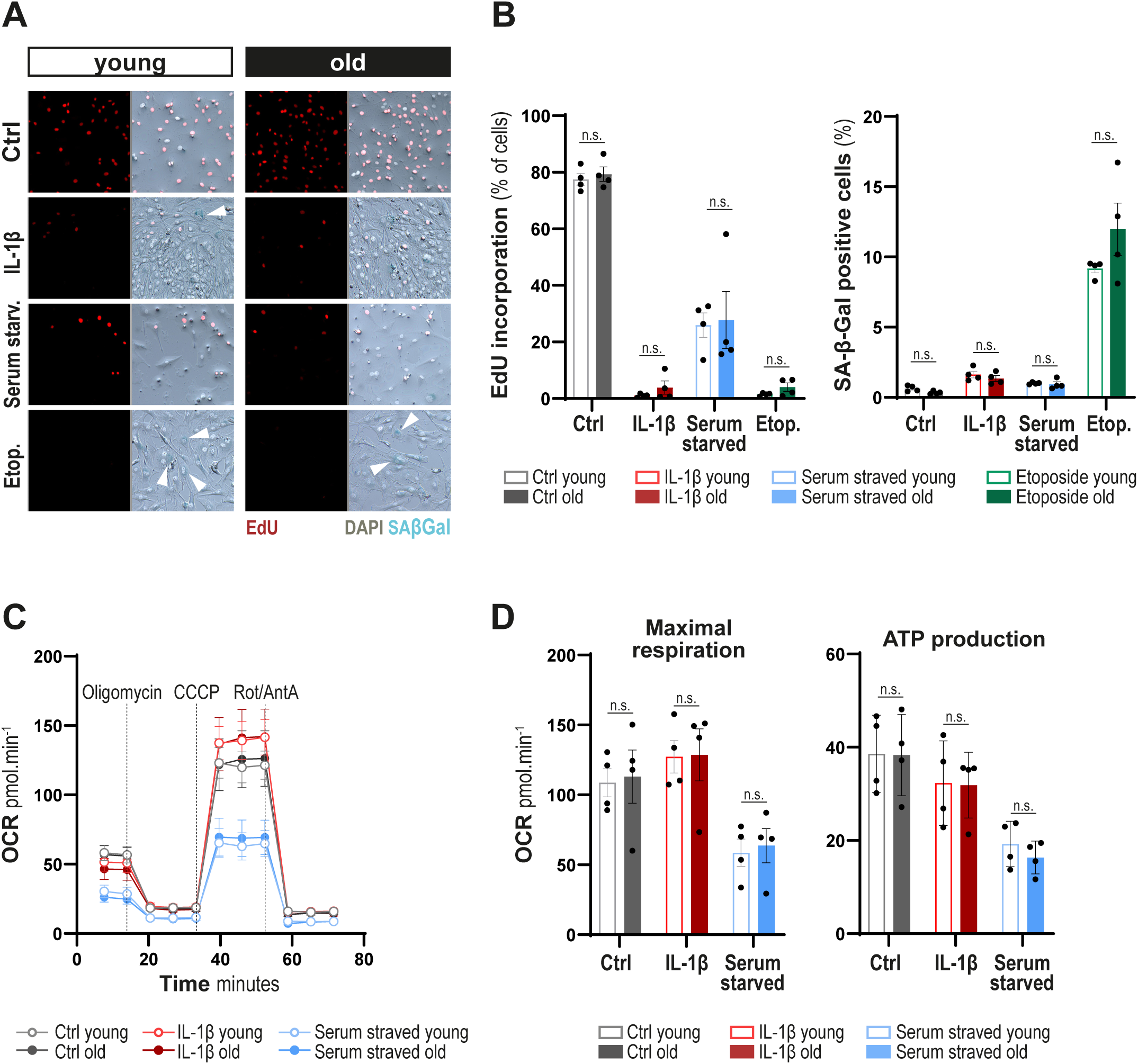
NPC from young and aged ovine disc share common senescence and mitochondrial energetic responses to stress-related stimuli. **A.** Co-immunofluorescence of EdU incorporation, an indicator of cell proliferation, and SA-β-Gal staining, indicating high lysosomal content and potential marker of cellular senescence. DAPI is used as a nuclear marker. Arrows highlight SA-β-Gal positive blue staining. **B.** Quantification of % of NPC from young and old animals positive for EdU incorporation and SA-β-Gal staining. **C.** Functional investigation of mitochondrial energetic profiles changes by Seahorse® evaluating Oxygen Consumption Rate (OCR) measurements using the mitostress test assay in NPC from young and old animals. **D.** Quantification of maximal respiration and ATP production. ns: not significant, p-values were calculated using the Kruskal-Wallis test, and Dunn’s post-hoc test; N=4 animals per group.

In control conditions, NPC from both young and old animals were proliferating similarly, with around 80% of them incorporating EdU overnight, and only marginally expressing SA-β-Gal (**Figure 4A, B, Supplementary Figure S5B**). All treatments greatly restricted cell proliferation, down below 40% EdU incorporation with serum starvation and even under 10% with IL-1β or etoposide treatment. Etoposide was the only treatment increasing SA-β-Gal expression up to 10% of stained cells. However, no difference in cell state were observed between cells from young and old animals, at baseline and after treatments.

Accordingly, NPC from young and aged animals exhibited similar mitochondrial energetic profiles in basal conditions and in responses to either IL-1β or serum starvation (**Figure 4C, D, Supplementary Figure S5C**). This included comparable maximal respiration and ATP production for NPC from young and old animals at baseline, and the same increase of maximal respiration and decrease of ATP production in IL-1β-treated condition. Lastly, serum starvation reduced maximal respiration and ATP production similarly in young and old NPC.

Therefore, these functional experiments highlighted that NPC derived from mildly degenerated discs remain responsive to inflammatory, senescence-inducing, and metabolic stress stimuli, similarly to NPC derived from healthy young IVD.

## 4. Discussion

The sheep model is already widely used and recognized for ex vivo and in vivo studies of IVDD [9], positioning ovine-derived cell culture models as particularly valuable tools to bridge basic and translational research. Considering that its naturally occurring degenerative process resembles the human one, cellular models from sheep are more likely to mimic what could be observed in human cells. Moreover, the availability of animals across a wide age range offers the opportunity to investigate developmental, maturational, and degenerative processes. However, this advantage comes with significant limitations. A key limitation of our study is the exclusive use of female animals. Livestock management practices primarily dictated this choice: in farming, male lambs are generally slaughtered for meat, whereas females are retained for breeding and milk production and can reach advanced ages (6–7 years). As a result, aged males are rarely available, and using females across all age groups avoids introducing a sex bias between cohorts. While sex-specific differences in IVDD have not been clearly established in sheep, they are well documented in humans. Biological factors such as hormones, immunity, behaviour, and gene expression contribute to sex-related differences in chronic diseases. Importantly, low back pain, a clinical manifestation of IVDD, shows consistently higher prevalence and incidence in women than in men across all age groups, with even more pronounced differences in older populations [31]. Therefore, although driven by practical considerations, the exclusive use of female sheep in this study may also increase the translational relevance of the model. Finally, this approach is consistent with the 3R’s principle of reduction, as we routinely obtain female animals from local farms, avoiding the need to breed males for research purposes specifically.

A clear strength of the ovine model lies in the anatomical size of the IVD close to the human ones, allowing for macroscopically distinguishing and dissecting the AF from the NP. This facilitates the independent isolation and culture of the two main disc cell types, a significant technical advantage over smaller animal models where such dissection is challenging and imprecise. In this study, we efficiently isolated and expanded AFC and NPC from all individuals across both age groups. AFC and NPC retained somewhat distinct transcriptomic signatures in vitro that were more marked in cells isolated from older animals, likely reflecting full maturation of the IVD tissue. Notably, AFC, often understudied in the context of IVDD, were readily available and displayed age-related transcriptomic features, supporting their inclusion in studies investigating disc degeneration and in the preclinical evaluation of biomaterials, such as those designed for herniated disc repair [32].

In addition to the technical and logistical burden that is represented by IVDs’ isolation from these large animals, the main drawback highlighted here is the pronounced inter-individual variability, illustrated by the absence of clear clustering on RNA-seq PCA plot. While this mirrors the biological heterogeneity seen in human populations and may thus increase translational relevance compared to studies using inbred rodent models or cell lines, it presents a challenge for statistical analyses and experimental reproducibility. The use of bioinformatics tools built around human gene sets further complicates interpretation, as species-specific differences in gene annotation and pathway mapping can introduce biases or lead to incomplete pathway inference. Nevertheless, the cross-species comparison of our data with a human dataset allowed us to confirm that mostly the “fibrotic” and, to a certain extent, the “progenitor” subsets of IVD cells seem to grow in vitro, whether they are the only ones able to survive and proliferate, or whether they are cellular states converging in culture due to dedifferentiation [33]. Future studies on these models should thus focus on the potential role of these specific resilient cell sub-populations in IVD degeneration and aim to modulate their phenotype. Importantly, linking the aging signature of sheep cells with data from human samples reinforces the expected similarity between ovine and human IVDD.

In the functional assays, sheep NP cells were compatible with all experimental technical approaches used with minimal adjustment to standard protocols. We used commercially available ovine recombinant IL-1β, but cells were also responsive to the human recombinant cytokine (data not shown). However, a notable limit of this model may come with the use of antibody-based approaches for which anti-sheep protein specificity may often be unproven or unavailable. Cells from young and old animals showed similar mitochondrial energetic profile and predisposition to senescence, independently of the treatment tested. Although surprising considering the differences observed at the transcriptomic level, these results could be partially explained by the use of cells with higher passage for functional assays, as we needed sufficient numbers for all analytical techniques and experimental conditions. On the bright side, these findings show that NPC derived from mildly degenerated discs remain viable and thus potentially responsive to regenerative stimuli in vivo. However, this functional result contrasts with studies in rabbits, where age-associated mitochondrial energetic dysfunction and increased senescence were reported in cultured IVD cells [34]. The discrepancy may be attributable to species-specific differences in disc biology; if rabbits have been shown to develop spontaneous IVDD [35], they also seem to maintain notochordal cells throughout adulthood, a major difference compared to sheep [11]. This maintenance of a progenitor cell pool could explain different aging processes, IVDD mechanisms and regenerative capacities between those species.

Altogether, our findings support the use of sheep-derived NP and AF cells as relevant in vitro models of early-stage IVDD. While cells from animals of all ages are suitable for experimentation, the focus should be placed on the biological stimulus rather than the donor age, with the understanding that aging markers may be subtle or absent with only a subset of cells surviving in vitro. Several strategies could enhance the physiological relevance of these in vitro models to recapitulate cellular states lost with initial cell isolation and expansion, including culture in three-dimensional spheroids [36], encapsulation in hydrogels mimicking disc matrix properties [37,38], or maintenance in hyperosmolar [39] or hypoxic conditions [40], which could better replicate the native disc microenvironment.

In conclusion, our study demonstrates that ovine NPC and AFC represent a promising model for the study of mechanisms implicated in disc aging and degeneration, and for the early evaluation of potential regenerative therapies, while suffering of the same limitations as any cell culture model. Similarities found with the transcriptomic signature of human cells reinforce the translational relevance of the ovine model from in vitro to in vivo experiments.

## List of abbreviations

AF: Annulus Fibrosus
AFC: Annulus Fibrosus Cells
CEP: Cartilaginous Endplate
GSEA: Geneset enrichment analysis
IVD: Intervertebral disc
IVDD: Intervertebral disc degeneration
LBP: Low back pain
NES: Normalized Enrichment Scores
NP: Nucleus Pulposus
NPC: Nucleus Pulposus Cells
OCR: Oxygen Consumption Rate
OXPHOS: Oxidative phosphorylation

## Declarations

### Ethics

The animals used in this study were housed in accordance with good practices and animal welfare standards, under the supervision of veterinarians. Euthanasia was performed according to good practices and legislation (European Directive 2010/63/EU) in the accredited Centre of Research and Pre-clinical Investigations (CRIP) at the ONIRIS-National Veterinary School of Nantes. All efforts were made to minimize the number of animals used and their suffering.

### Availability of data and materials

The raw and processed transcriptomic data that support the findings of this study have been deposited in NCBI’s Gene Expression Omnibus and are accessible through GEO Series accession number **GSE302551**. The complete bioanalysis pipeline is available at https://gitlab.univ-nantes.fr/guihomics/ovine_IVD_bulkRNAseq. Other raw data and materials are available from the corresponding author on reasonable request.

### Authors’ contributions

RG, JG and CLV conceived and designed the study. PH, LD, EC and MF performed the formal analyses. JG, CLV, FB and RG acquired funding. PH, LD, EC, BH and FE carried out investigations. PH, LD, RG, CV, MF and FB developed the methodology. PH and RG drafted the original manuscript. All authors read, amended, and approved the final version of the manuscript.

## Acknowledgments

The authors thank O. Gauthier, A. Lafragette, P. Roy, S. Madec, D. Rouleau, P. Monmousseau from the Center for Research and Preclinical Investigation (C.R.I.P.; Oniris Nantes Atlantic College of Veterinary Medicine, Food Science and Engineering, Nantes, France) for their professionalism and precious help in this study.

The authors thank C. Pecqueur (CRCI2NA, Nantes Université, INSERM U1307, CNRS 6075, Nantes, France) for her help with Seahorse technology, with financial support from ITMO Cancer of Aviesan within the framework of the 2021-2030 Cancer Control Strategy.

The authors acknowledge the SC3M platform from the Inserm/NU/ONIRIS UMR1229 RMeS Laboratory and SFR Bonamy. They also acknowledge the MicroPICell core facility (SFR Bonamy, BioCore, Inserm UMS 016, CNRS UAR 3556, Nantes, France), member of the Scientific Interest Group (GIS) Biogenouest, IBISA, and the national infrastructure France-Bioimaging supported by the French national research agency (ANR-10-INBS-04).

The authors thank the Genomics Core Facility GenoA, member of Biogenouest and France Genomique, and the Bioinformatics Core Facility BiRD, member of Biogenouest and Institut Français de Bioinformatique (ANR-11-INBS-0013) for the use of their resources and their technical support.

Schematics were created with Biorender.com.

---

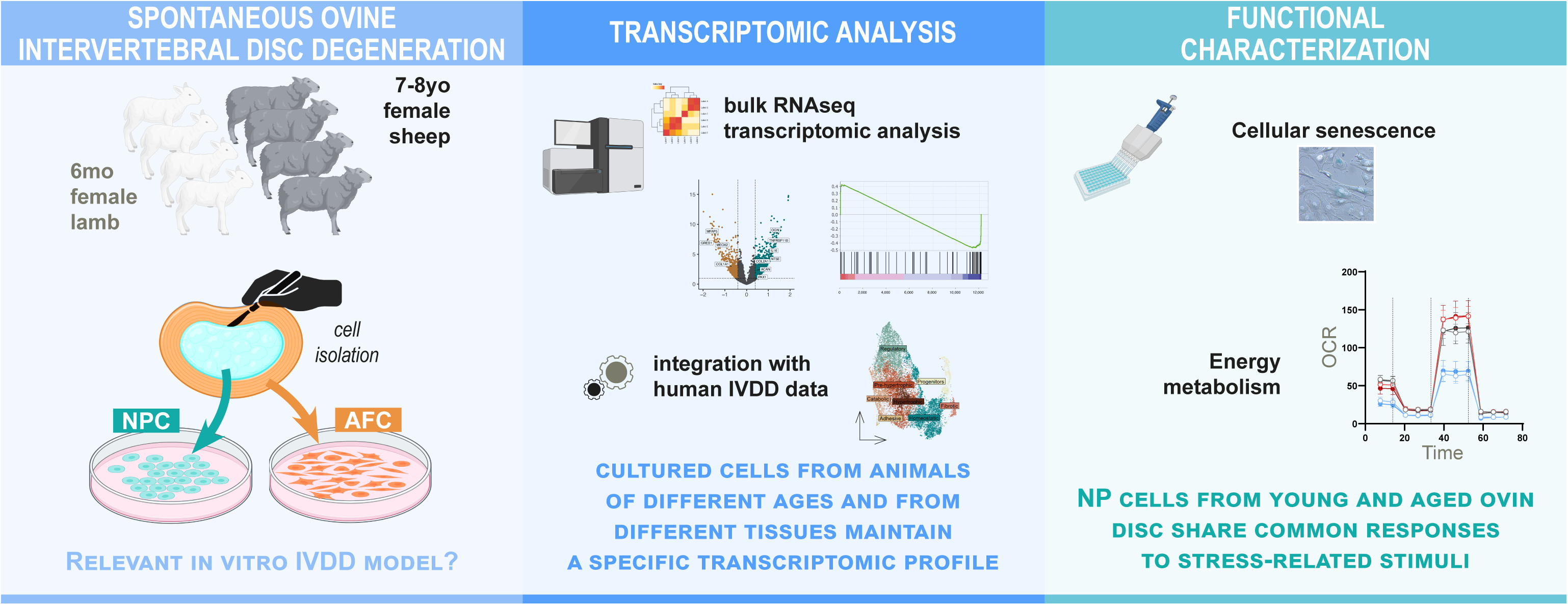

