## Supplementary material for "Transcriptomic and functional comparison of cells isolated from healthy and degenerated ovine intervertebral discs"

#### Supplementary Methods

#### Supplementary Figures

#### Supplementary Tables legends

### Supplementary Methods

#### Quantitative RT-PCR Analysis of IL-6 and MMP3 Expression in NP Cells

NP cells at P3 were seeded in 6-well plates at 100,000 cells per well. After 3 days, the culture medium was changed after a PBS wash to (1) complete culture medium, (2) complete culture medium supplemented with ovine IL-1β (10ng/mL), or (3) serum-free culture medium. After 24h, total RNA was extracted using the Nucleospin RNA XS kit (Macherey-Nagel, 740902) according to the manufacturer’s instructions. RNA yield was measured using NanoDrop 1000 Spectrophotometer (Thermo Scientific). Reverse transcription was performed using the Verso cDNA Synthesis Kit (Thermo Scientific, AB1453B) with a mix of Anchored Oligo(dT) and Random Hexamers. Real-time polymerase chain reaction (PCR) was performed using specific *ovis aries* primers (5’ -> 3’: IL6_Forward: CCTGTCCACTGGGCACATAA; IL6_Reverse: GTTCAAGCCGCATAGCCATT; MMP3_Fw: GTGTGATCCTGCCTTGTCCT; MMP3_Rev: CCGCCAAAAATGTCTGCCTTT; housekeeping gene: GAPDH_Fw: TCCTGCCCCACCTCCACCAC; GAPDH_Rev: GGGCTCCCTAAGCCCCTCCC) with SYBR Select Master Mix (Applied Biosystems, 4472908) on the CFX96 Touch Real-time PCR Detection System (Bio-Rad).

### Supplementary Figures

#### Supplementary Figure S1

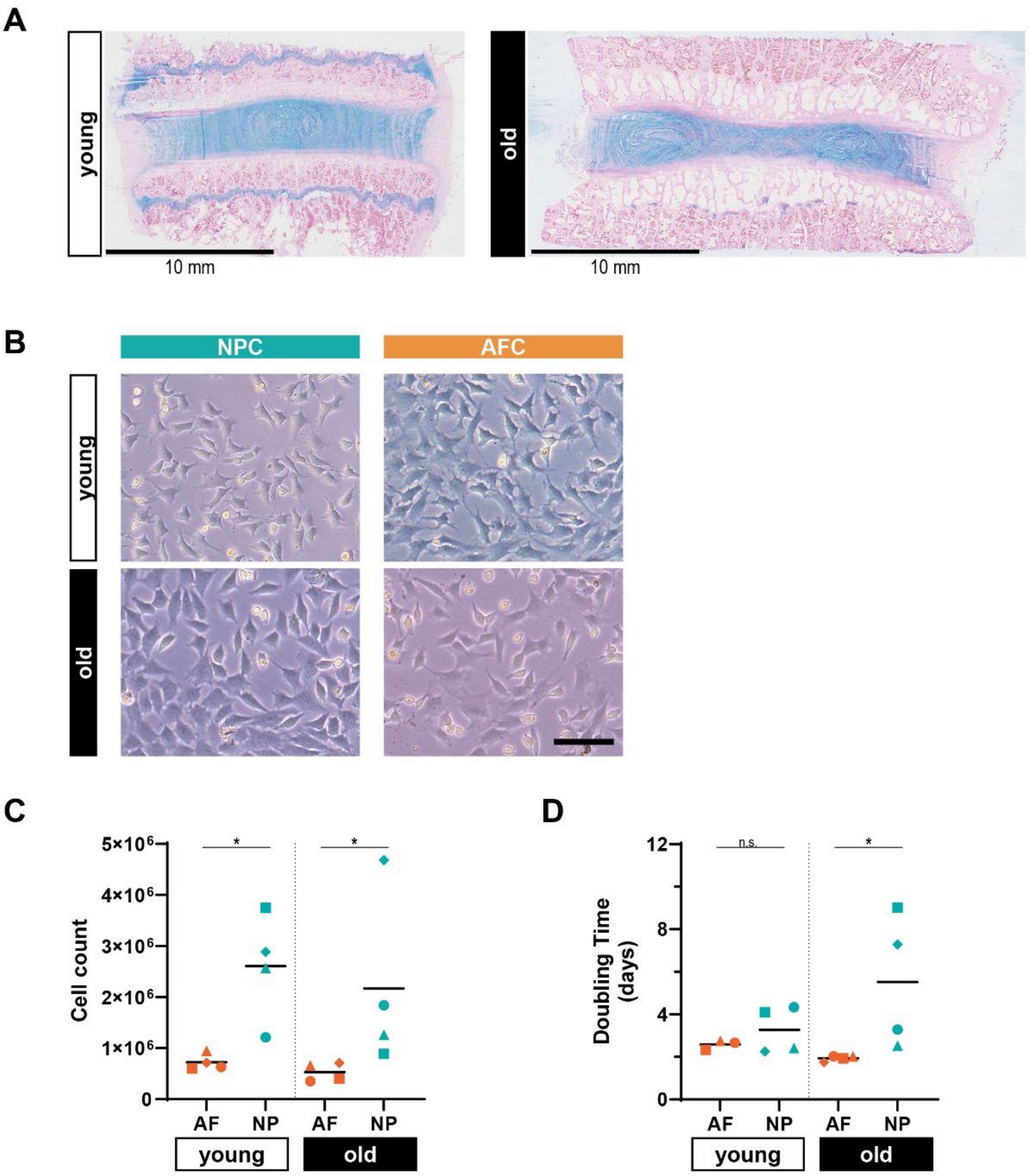


**Supplementary Figure S1 – Relative to Main Figure 1. A.** Representative Alcian Blue staining images. Scale bar: 10 mm. **B.** Cell morphology at passage P1. Scale bar: 100µm. **C.** Cell count after cell isolation. **D.** Cellular doubling time between passage P0 and P1. n.s.: not significant; *: $p<0.05$. p-values were calculated using Mann Whitney test; N=4 young and N=4 old animals.

**Supplementary Figure S2**

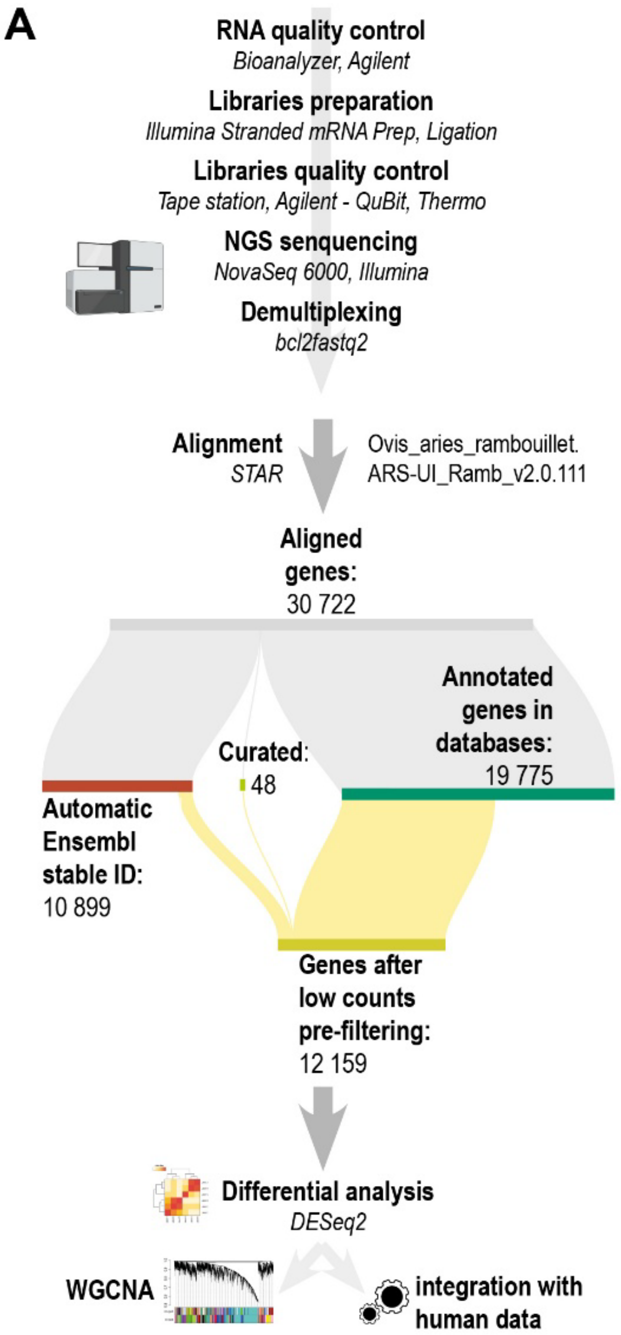


**Supplementary Figure S2. Schematic bioinformatic analysis pipeline. A.** From primary to secondary analysis pipeline.

**Supplementary Figure S3**

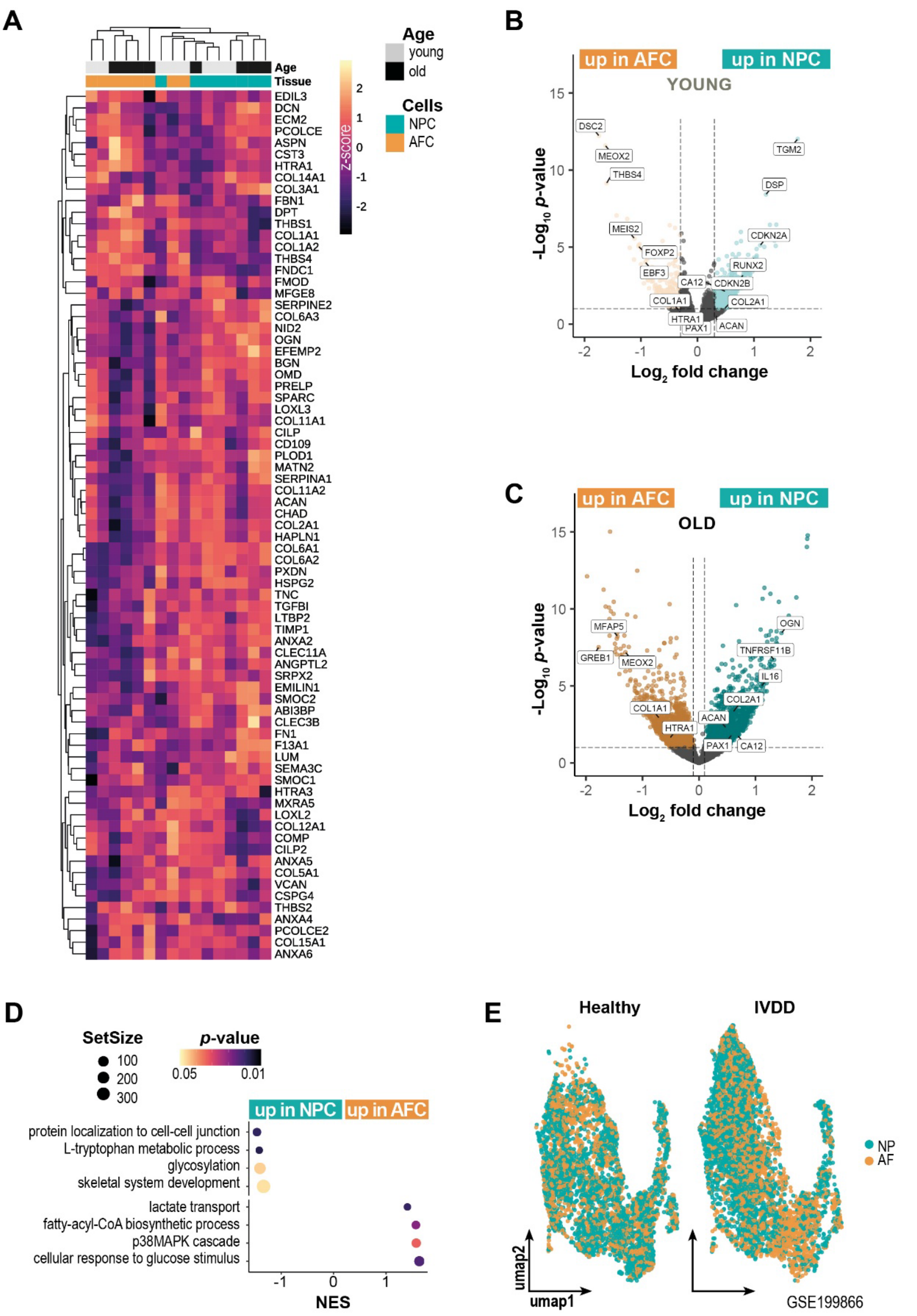


**Supplementary Figure S3 – *Relative to Main Figure 2.* A.** Heatmap showing sample clustering based on matrix-related transcripts expression from DIPPER analysis (Tam et al.

2020). **A**. Volcano plot of differential gene expression in NPC versus AFC from young and **B.** old animals using the Wald test. **D.** Geneset Enrichment Analysis (GSEA) AFC versus NPC samples using the permutation test. **E.** UMAP of scRNA-seq human dataset (GSE199866) split between one healthy and one IVDD intervertebral disc, cells from NP tissue are highlighted in turquoise and from AF tissue in orange.

**Supplementary Figure S4**

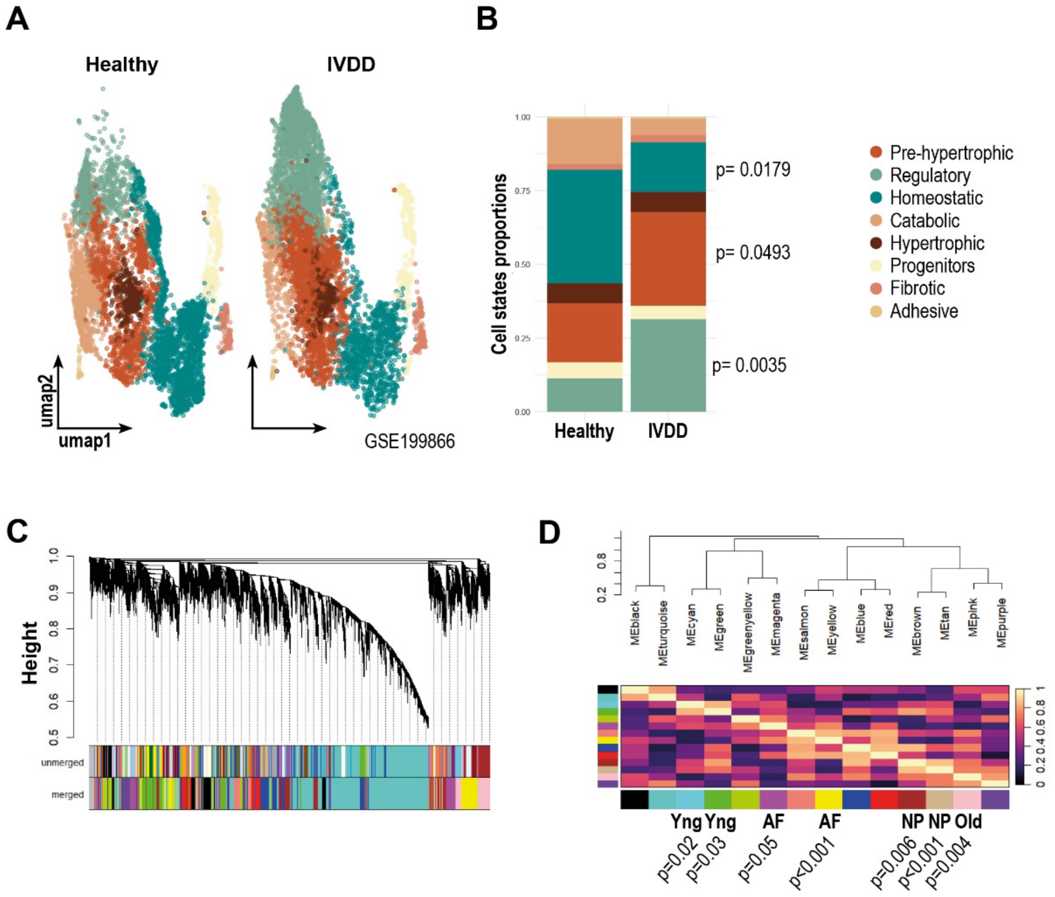


**Supplementary Figure S4 – *Relative to Main Figure 3*. A.** UMAP of scRNA-seq human dataset (GSE199866) used as basis for bulk RNA-seq deconvolution combining AFC and NPC transcriptome, split between one healthy and one degenerated intervertebral disc. Cellular states were annotated using literature consensus. **B.** Proportion of the different cellular states present in healthy and IVDD condition. **C.** Dendrogram plot showing hierarchical clustering of the 12159 genes in 14 modules after merging, a different colour represents each module. **D.** Dendrogram and heatmap generated with the module eigengenes for the 14 modules identified. Significant association between modules and origin of the cells are highlighted below (Yng: young). p-values were calculated with the Student asymptotic p-value for correlation.

**Supplementary Figure S5**

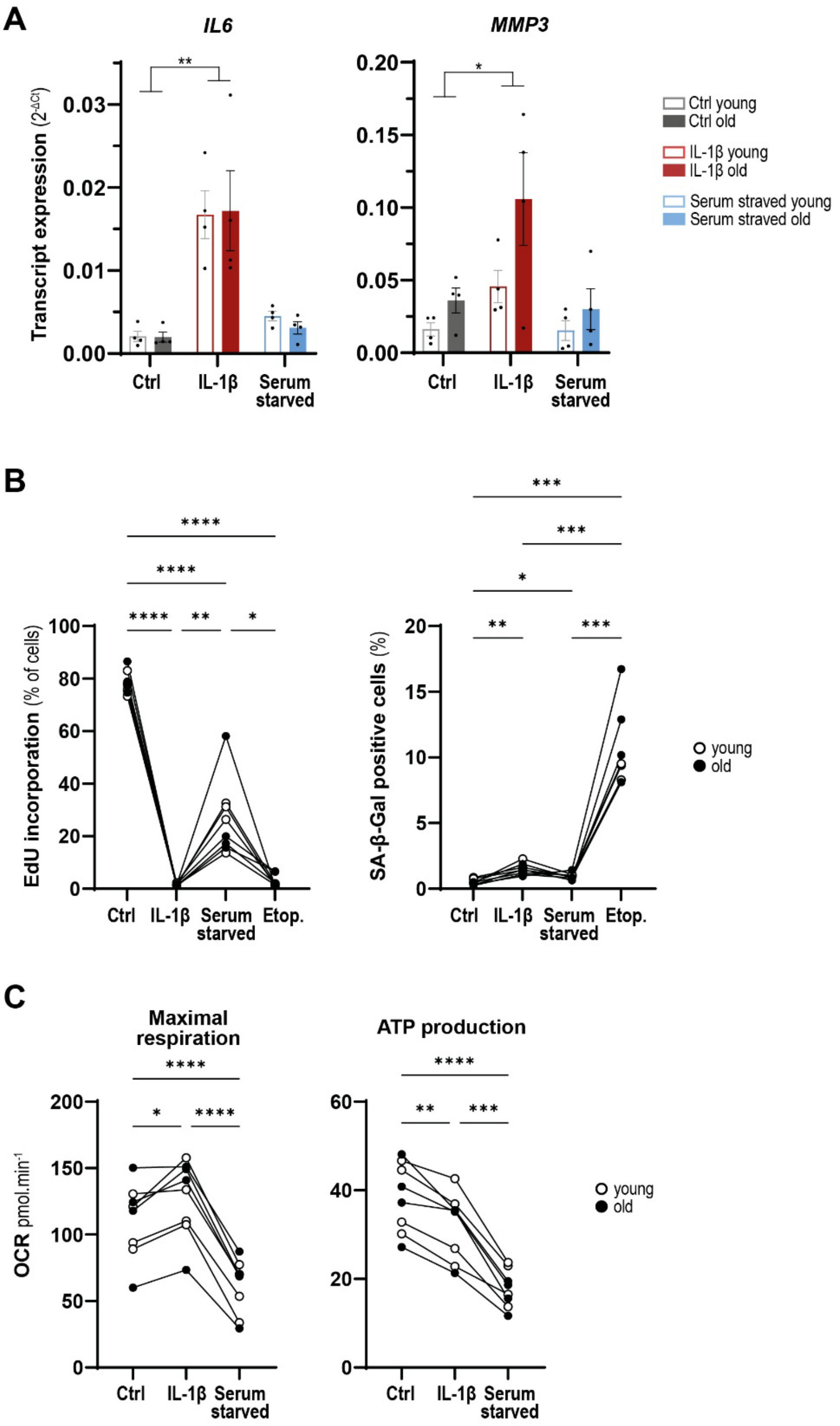


**Supplementary Figure S5 –** ***Relative to Main Figure 4*****. NPC from young and aged ovine disc share common senescence and mitochondrial energetic responses to stress-related stimuli. A.** IL6 and MMP3 transcript expression by RT-qPCR of NPC from young and old animals following 24h of IL-1 β treatment or serum starvation. **B.** Quantification of % of NPC from young and old animals positive for EdU incorporation and SA-β-Gal staining. **C.**

Quantification of maximal respiration and ATP production. *:$p<0.05$; **:$p<0.01$; ***:$p<0.001$; ****:$p<0.0001$. p-values were calculated using the Repeated Measures ANOVA test, and Tukey's post-hoc test; N=8 animals.

#### Supplementary Tables legends

**Supplementary Table 1. Differential Expression Results between NPC from young and old animals.** Standard differential expression analysis results using DESeq2 pipeline. baseMean= mean of normalized read count across all samples. log2 fold change= log2 Fold change between groups, positive indicates upregulated in NPC isolated from old animals, padj= p-value adjusted for multiple testing using Benjamini-Hojberg correction.

**Supplementary Table 2. Differential Expression Results between AFC from young and old animals.** Standard differential expression analysis results using DESeq2 pipeline. baseMean= mean of normalized read count across all samples. log2 fold change= log2 Fold change between groups, positive indicates upregulated in AFC isolated from old animals, padj= p-value adjusted for multiple testing using Benjamini-Hojberg correction.

**Supplementary Table 3. Differential Expression Results between NPC and AFC isolated from young animals.** Standard differential expression analysis results using DESeq2 pipeline. baseMean= mean of normalized read count across all samples. log2 fold change= log2 Fold change between groups, positive indicates upregulated in NPC, padj= p-value adjusted for multiple testing using Benjamini-Hojberg correction.

**Supplementary Table 3. Differential Expression Results between NPC and AFC isolated from old animals.** Standard differential expression analysis results using DESeq2 pipeline. baseMean= mean of normalized read count across all samples. log2 fold change= log2 Fold change between groups, positive indicates upregulated in NPC, padj= p-value adjusted for multiple testing using Benjamini-Hojberg correction.

**Supplementary Table 5. List of the top-30 genes associated with WGCNA modules.** List of the 30 genes most associated with the different modules identified by WGCNA, ranked by importance in the module. The modules included are the modules associated with AFC (Yellow and Magenta modules), with NPC (Tan and Brown modules), with young and (Cyan and Green modules) and with old age (Pink module).
